# Nuclear Myosin 1 links genomic architecture to adipose tissue remodeling, metabolic inflammation and obesity in mice

**DOI:** 10.1101/2025.06.30.662298

**Authors:** Samira Khalaji, Tomas Venit, Zuzana Lukacova, Valentina Fambri, Rahul Shrestha, Sachin Kaluarachchi, Maylis Boitet, Maud Fagny, Giuseppe Saldi, Piergiorgio Percipalle

## Abstract

During adipogenesis, a metabolic shift from oxidative phosphorylation (OXPHOS) to aerobic glycolysis enables preadipocytes to meet the biosynthetic and energetic demands of differentiation. Nuclear myosin 1 (NM1), a chromatin-associated actomyosin motor that regulates transcription and chromatin accessibility, is essential for maintaining OXPHOS. Here, we identify NM1 as a key regulator of adipocyte differentiation and adipose tissue homeostasis. Integration of ATAC-seq and RNA-seq in NM1-deficient mouse embryonic fibroblasts revealed coordinated dysregulation of adipogenic genes (*Insig1, Lipg, Fat1*) and altered enhancer accessibility near *Klf6, Foxo3, Smad5*, and *Gata4*. NM1 knockout mesenchymal stem cells showed impaired adipogenic differentiation despite adipocyte hypertrophy. In vivo, NM1-deficient mice developed age-dependent visceral obesity with transcriptional reprogramming in white adipose tissue (WAT), including downregulation of adipogenesis and mitochondrial pathways, and activation of IFNG-, IL33-, and TNF-driven inflammation. Cross-species analysis revealed overlap with *MYO1C*-centered regulatory modules in human adipose tissue, implicating NM1/*<u>MYO1C</u>* in conserved chromatin-based control of adipose biology

## Introduction

Mitochondria are the principal source of Adenosine triphosphate (ATP) through oxidative phosphorylation (OXPHOS) and function as central regulators of intracellular calcium signaling, metabolite sensing, and epigenetic modification through TCA cycle intermediates^1,2^. Their biogenesis and roles are highly dynamic and adapt to the metabolic demands of specific cellular states. This is particularly important for cells undergoing commitment to specific lineages when there is a need to increase OXPHOS activity while decreasing glycolytic flux, a metabolic transition essential for lineage fidelity and organ function^3,4^. During adipogenesis, however, this classical model is reversed in early phases. Committed preadipocytes undergo a shift from OXPHOS to aerobic glycolysis, a form of metabolic reprogramming that prioritizes rapid ATP generation and the production of biosynthetic intermediates. Although glycolysis is less efficient than OXPHOS in terms of ATP yield, it provides critical substrates for lipid and nucleotide biosynthesis, fueling cell growth and lipid droplet formation. Additionally, the reduction in mitochondrial activity during this shift minimizes reactive oxygen species (ROS) generation, protecting differentiating cells from oxidative stress^5^. As differentiation proceeds, mitochondrial biogenesis is reactivated, and OXPHOS capacity increases to support the energy demands of mature adipocytes. This metabolic remodeling is tightly linked to the transcriptional activation of adipogenic regulators such as *PPARγ* and *C/EBPα*. Furthermore, hypoxia-inducible factors (HIFs) and other signaling molecules activated during glycolytic metabolism may contribute to adipocyte phenotype commitment^6^. Overall, the switch to aerobic glycolysis supports the anabolic and redox needs of adipogenesis, ensuring proper adipocyte development and function. Disruptions in this metabolic adaptation can impair adipocyte formation and contribute to metabolic disorders such as obesity and insulin resistance. Therefore, understanding this shift is crucial for targeting adipose tissue dysfunction in metabolic diseases. During adipogenesis, MSCs differentiate into adipocytes and this process is fundamental to the development of WAT, a central endocrine organ. WAT governs energy storage and systemic metabolic homeostasis and participates in immune modulation, reproductive signaling, and angiogenesis^7,8^. Dysregulated adipose tissue expansion contributes to obesity, which is associated with adipocyte hypertrophy, chronic low-grade inflammation, and altered lipid metabolism. These features are mechanistically linked to metabolic pathologies including type 2 diabetes and cardiovascular disease^9^. However, the transcriptional and epigenetic regulators that coordinate adipogenesis and maintain adipose tissue function remain incompletely understood. Recently, we demonstrated that nuclear-encoded mitochondrial transcription factors are critically regulated by NM1^10^. NM1 deletion induces a metabolic switch from OXPHOS to aerobic glycolysis, a hallmark of metabolically reprogrammed or transformed cells, suggesting a tumor suppressor-like role for NM1, potentially through p53 signaling^11^. In the cell nucleus, cytoskeletal proteins including β-actin and several myosin species have emerged as key regulators of genome organization and gene expression. Both β-actin and NM1 interact with all three nuclear RNA polymerases to modulate transcriptional output^12,13,13–15^ . NM1, the well-characterized isoform b of *MYO1C*, acts as a component of the B-WICH chromatin remodeling complex, facilitating transcription by RNA polymerases I and II^16,17^. NM1 functions in this complex by promoting the nucleosome repositioning activity of B-WICH and recruitment of histone acetyl-transferases (HATs) and methyltransferases (HMTs), including *PCAF* and *Set1B*. This promotes histone acetylation and methylation at gene promoters and enhances chromatin accessibility^16,17^. Remarkably, loss of NM1 led to altered key metabolic pathways related to mitochondrial function, including cellular signaling cascades, nutrient sensing mechanisms, and epigenetic regulation mediated by TCA cycle metabolites^11,18,19^. Considering its role in transcription and chromatin regulation, NM1 represents a unique factor at the interface of transcription, epigenetics, and cellular metabolism. Here, we hypothesized that NM1 serves as a key regulatory node in adipogenesis. Using a machine learning approach, we began by establishing a predictive model based on single cell RNA-seq and ATAC-seq datasets to infer transcription factor activity and predict gene expression responses to NM1 deletion. We next integrated ATAC-seq and RNA-seq data from WT and NM1 KO MEFs^20^ to establish a potential involvement of NM1 in key cellular processes (Figure 1A). Results from these experiments indicate that NM1 regulates key adipogenic factors such as *Insig1*, *Insig2* and *Lipg* through a chromatin-based mechanism. To test this further, we used a loss of function mouse model and performed a comprehensive analysis combining ex vivo adipocyte differentiation from MSCs, transcriptional profiling of VAT, and cross-species eQTL network analysis. In addition, Ingenuity Pathway Analysis (IPA) of RNA-seq data from VAT revealed dysregulation of key metabolic and inflammatory signaling pathways, further implicating NM1 in adipose tissue function. Our findings reveal a multifaceted role for NM1 in coordinating adipocyte differentiation, mitochondrial remodeling, and adipose tissue inflammation. Together, these findings point to NM1 as an essential chromatin regulator for maintaining adipose tissue homeostasis and suggest it may play a role in the molecular mechanisms underlying obesity.

**Figure 1.**
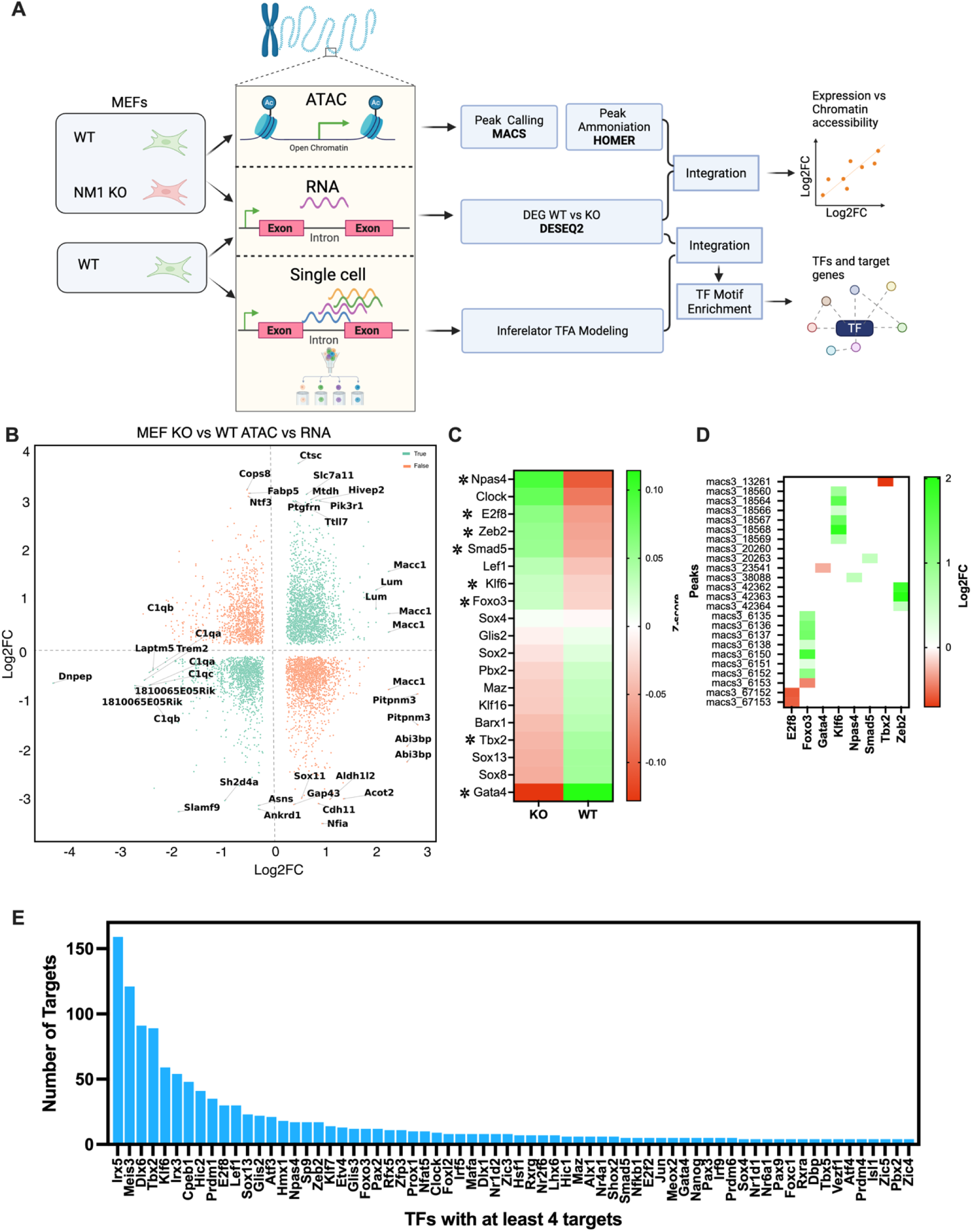
NM1 regulates transcription factor networks through integrated chromatin accessibility and transcriptomic modeling in MEFs. (A) Schematic overview of the integrative pipeline combining ATAC-seq, bulk RNA-seq, and single-cell RNA-seq from WT and NM1 KO MEFs. ATAC-seq peaks were identified using MACS and annotated with HOMER, while differentially expressed genes (DEGs) were detected using DESeq2. The datasets were integrated to identify genes with concordant chromatin and expression changes and to infer TF–target gene relationships using the Inferelator framework and motif enrichment (right panels). Visualization of transcription factors identified from the inferred TF–target gene network shown in Supplementary Figure 1, based on integrated RNA-seq, ATAC-seq, and single-cell data (B) Scatter plot showing log2 fold changes in gene expression (RNA-seq) versus chromatin accessibility (ATAC-seq) between WT and KO MEFs. Genes with significant concordant changes in both are highlighted in orange. (C) Heatmap of RNA-seq expression values (z-score normalized) for the top 19 transcription factors identified in the inferred regulatory network. Asterisks (*) indicate transcription factors whose chromatin accessibility is significantly altered in KO versus WT. (D) Chromatin accessibility (log2FC) at MACS3-called ATAC-seq peaks associated with the 8 transcription factors from panel C showing significantly altered accessibility. Positive values indicate increased accessibility in KO cells; negative values indicate decreased accessibility. (E) Bar plot of transcription factors from the inferred network regulating at least 4 target genes. The number of downstream targets is plotted for each TF, ranked from highest to lowest.

## Results

### NM1 regulates key adipogenic factors through a chromatin-based mechanism

Given NM1’s established role in chromatin remodeling and transcriptional regulation, we studied whether NM1 serves as a key regulatory node in adipogenesis, controlling adipogenic transcription factors through a chromatin-based mechanism. To this aim, we integrated differential gene expression and chromatin accessibility data previously generated in our lab in WT and NM1 KO MEF^10,11^ and established a predictive regulatory model that integrates chromatin accessibility with single-cell RNA-seq data to infer transcription factor activity and network structure. Genes exhibiting significant changes in expression (adjusted p-value ≤ 0.05 and |logFC| ≥ 0.25) were compared to regions of differential accessibility (FDR ≤ 0.05 and |logFC| ≥ 0.1). While a proportion of differentially expressed genes did not exhibit corresponding changes in chromatin accessibility, suggesting indirect or post transcriptional effects, a distinct subset showed coordinated alterations in both gene expression and chromatin state (orange dots in Figure 1B). Concordant alterations in both expression and accessibility, suggests that changes in chromatin accessibility may directly influence their transcriptional regulation in the absence of NM1 (Figure 1B). Among these, several genes were functionally linked to adipogenesis or lipid metabolism (supplementary table2). For example, *Insig1* and *Insig2*, both negative regulators of the SREBP pathway, showed significantly decreased expression and chromatin accessibility in NM1 KO cells^21,22^. *Lipg* and *Mylip*, involved in lipoprotein processing and lipid homeostasis, and *Fat1*, associated with adipocyte expansion, were also repressed and less accessible in KO cells^23–26^. Additional genes such as *Acsl4*, *Cd9*, *Lpin1*, *Fndc5*, and *Wnt11* showed coordinated downregulation and loss of chromatin accessibility, further supporting NM1’s role in regulating lipid metabolism and adipogenic potential^27–30^. These findings suggest that NM1 may control adipogenic capacity in part through direct regulation of lipid-handling genes at the chromatin level. To further investigate this correlation, we created a gene regulatory network (GRN) by integrating chromatin accessibility data to identify transcription factor binding motifs and link them to nearby genes (Supplementary Figure 1B), using the inferelator-prior tool with the HOCOMOCO TF database. To simulate NM1 knockout expression, transcription factor activity (TFA) from the bulk RNA KO experiment was multiplied by the wild-type regulatory network, and the simulated expression values were compared to the observed measurements (R² = 0.65) to evaluate network accuracy (Supplementary Figure 1A). Motif enrichment analysis using FIMO and the HOCOMOCO database identified candidate TF binding sites within these accessible regions (Figure 1E). This analysis revealed 68 highly active TFs in WT MEFs, each regulating at least four target genes. From this list, we selected 19 transcription factors for further analysis based on their known or putative roles in adipogenesis and metabolic regulation. Many of these TFs, such as *Klf6*, *Foxo3*, *Smad5*, and *Gata4*, have been shown to regulate adipocyte differentiation, lipid metabolism, and mitochondrial function in adipose tissue^23–25^ . To refine the regulatory role of NM1 in transcription factor control, we compared chromatin accessibility (ATAC-seq) and expression (RNA-seq) of the top 19 TFs identified from our regulatory network model. Of these, 8 TFs exhibited significant differential accessibility in KO MEFs (Figure 1D, marked with asterisks in Figure 1C). *Foxo3*, *Klf6*, *Npas4*, and *Zeb2* showed both increased expression and chromatin accessibility in KO MEFs, suggesting NM1 may normally act to restrict their transcription through chromatin compaction. In contrast, *Gata4* and *Tbx2* displayed reduced expression and accessibility, indicating that NM1 is required to maintain an open chromatin configuration at these loci. *Smad5* presented increased accessibility but was transcriptionally downregulated, while *E2f8* showed the opposite pattern. These decoupled TFs suggest additional layers of transcriptional regulation that may involve NM1 independent mechanisms or control. All differentially accessible peaks were located in intergenic regions within 10–300 kb of the respective transcription start sites, suggesting a role for NM1 in enhancer-mediated transcriptional regulation. These findings highlight a dual role for NM1 in maintaining chromatin balance across key regulatory nodes during adipogenic differentiation. Further, using HiC-seq analysis we investigated if some of these transcription factors whose activity is affected through a chromatin-based mechanism in the absence of NM1 are also in chromatin compartments that undergo significant compartment switching. Although none of the 19 transcription factors in our network overlapped with Hi-C compartment switching regions, we identified several genes of interest among the switchers (Table 1). Notably, Ncoa2, a known coactivator of *PPARγ* and regulator of adipogenesis, transitioned between compartments in NM1 KO cells^26^. Other chromatin-associated or adipogenesis-linked genes, including *Cops5* and *Prdm14*^31,32^, also exhibited compartment switching, suggesting that NM1 may indirectly influence adipogenic programs through higher-order chromatin reorganization. These results altogether, underscore the importance of NM1 as a key transcriptional node in the regulatory landscape, where chromatin dynamics partially explains changes in transcription during adipogenesis.

**Table 1.** Genes exhibiting chromatin compartment switching in NM1 KO MEFs.

| **Gene** | **Compartment Switch**<br>**(WT → KO)** | **Functional Role** | **Biological Relevance** |
| --- | --- | --- | --- |
| **Ncoa2** | A → B | Nuclear coactivator of PPARγ | Adipogenesis, metabolic transcription |
| **Cops5** | A → B | COP9 signalosome subunit, chromatin remodeler | Gene expression, proteasomal regulation |
| **Prdm14** | B → A | Epigenetic regulator of pluripotency and fate | Stemness, chromatin organization |
| **Il18r1** | B → A | Interleukin-18 receptor | Inflammatory signaling in adipose tissue |
| **Il18rap** | B → A | IL-18 receptor accessory protein | Immune response modulation in VAT |

### NM1 deficiency leads to impaired adipogenesis *in vitro*

To investigate the role of NM1 in adipogenesis, MSCs isolated from WT and NM1 KO mice were differentiated into adipocytes over 20 days (Figure 2A). Morphological and quantitative analyses were performed at day 0 (pre-differentiation), day 5, day 7, day 10, day 15, and day 20. At day 0, MSCs from both genotypes exhibited similar spindle-shaped morphology without observable lipid accumulation. By day 5, lipid droplet formation had initiated, and by day 20, KO adipocytes displayed larger lipid droplets than WT, suggesting increased lipid storage capacity. Quantification of the adipocyte area supported these morphological observations (Figure 2C). At all the time points, KO adipocytes had significantly larger mean areas compared to WT. At day 20, KO adipocytes reached a mean area of 3583 æ165.1 μm² versus 2977 æ114.5 μm² in WT, representing a 20.4% increase. The most pronounced difference occurred on day 10, when KO adipocytes were 66.3% larger (2795 æ147.6 μm² vs. 1681 æ52.85 μm²). Standard deviation values were consistently higher in KO samples, indicating greater variability in adipocyte size. Despite the hypertrophy, KO cultures contained fewer adipocytes, suggesting impaired differentiation (Figure 2D).

**Figure 2.**
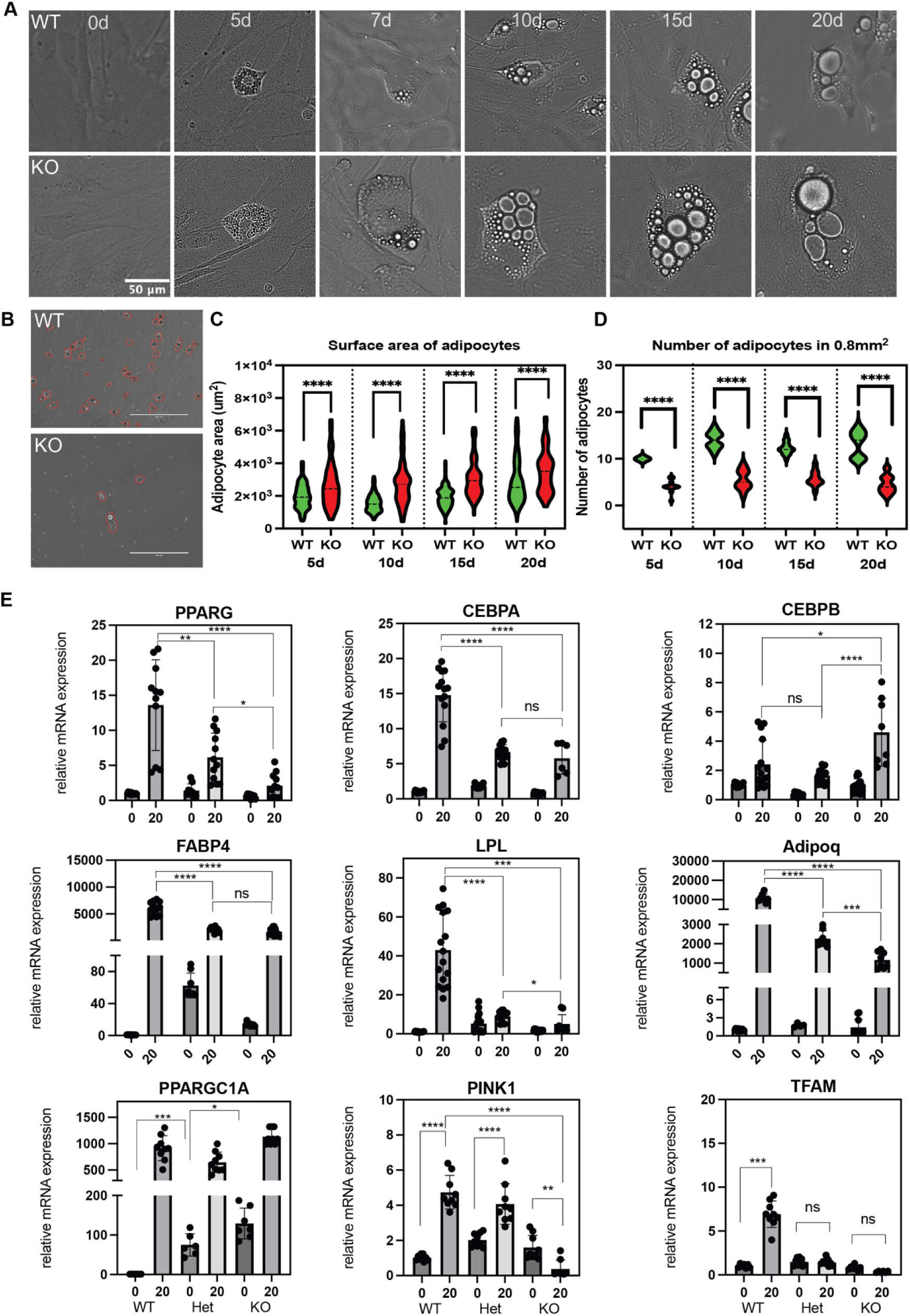
NM1 KO impairs adipogenic differentiation and alters adipocyte morphology and gene expression. (A)Time-course images of WT and NM1 KO MSC differentiation from day 0 to day 20. Scale bar: 50μm. (B) Representative images of WT and KO cultures at day 20, with adipocytes outlined in red for quantitative comparison of adipocyte number. Scale bars: 100 μm. (C) Violin plots quantifying adipocyte area (****p < 0.0001, unpaired t-test). (D) number of adipocytes per imagingarea ) at days 5, 10, 15, and 20. (****p < 0.0001, unpaired t-test). (E) Quantitative RT-PCR analysis of adipogenic markers (PPARG, CEBPA, CEBPB, FABP4, LPL, Adipoq) and mitochondrial regulators (PPARGC1A, PINK1, TFAM) in WT, heterozygous (HET), and KO MSCs at day 0 and day 20. Data represent mean ± SEM of at least three biological replicates; statistical significance was assessed using one-way ANOVA with post hoc tests (*p < 0.05; **p < 0.01; ***p < 0.001; ****p < 0.0001; ns, not significant).

We next quantified adipogenic markers expression in WT, HET (heterozygous), and KO MSCs before (day 0) and after differentiation (day 20) (Figure 2E). On day 20 post differentiation, *PPARγ* expression was highest in WT (13.61 æ6.47 fold), intermediate in HET (6.15 æ3.46), and lowest in KO (2.12 æ1.74). A similar trend was observed for *CEBPα*: 14.79 æ3.79 in WT, 6.68 æ1.05 in HET, and 5.78 æ2.26 in KO. In contrast, *CEBPβ* showed modestly higher expression in KO (4.61 æ 2.31) than in WT (2.42 æ1.65) and HET (1.65 æ0.49). Other markers, including *FABP*4, *LPL*, and *Adipoq*, followed the WT > HET > KO pattern. *FABP4*: 6017 æ1225 (WT), 2144 æ475.5 (HET), 1728 æ671.3 (KO); *LPL*: 43.04 æ18.43 (WT), 8.94 æ2.73 (HET), 5.05 æ4.79 (KO); Adipoq: 10888 æ2080 (WT), 2268 æ412.5 (HET), 1162 æ421.3 (KO). RT-qPCR analysis of mitochondrial regulators showed that *PPARGC1A* expression was elevated at baseline in KO (129.0 æ39.1-fold) and HET (74.9 æ28.2-fold) compared to WT (set to 1). Following differentiation, *PPARGC1A* was robustly upregulated in all genotypes: 915.0 æ238.8 (WT), 646.5 æ194.2 (HET), 1133.3 æ147.8 (KO). *TFAM* expression was similar at day 0 across genotypes, but post-differentiation induction occurred only in WT (6.92 æ1.50-fold), and was blunted in HET (1.43 æ0.39) and KO (0.35 æ 0.08). *PINK1* levels were slightly elevated at day 0 in HET and KO, but upon differentiation, only WT and HET showed strong induction. KO cells failed to increase *PINK1* (0.38 æ0.50-fold), suggesting impaired mitophagy.

Taken altogether, these results indicate that NM1 has a direct role in promoting efficient adipogenic commitment by regulating expression of adipogenic genes and mitochondrial adaptation during adipogenesis. Importantly, fewer but bigger adipocytes typically indicate that adipogenesis is impaired, and fat storage may be occurring via hypertrophy rather than healthy hyperplasia. This condition is associated with metabolic dysfunction and increased risk of obesity-related diseases.

### NM1 Knockout Results in Body Weight Gain and Enhanced Visceral Adipose Tissue Deposition

To find out if adipogenesis is dysregulated *in vivo* and leads to an obesity phenotype, we monitored NM1 KO and WT mice over time to assess body weight and adipose tissue accumulation. Upon dissection, the comparison of WT and NM1 KO mice showed visibly larger subcutaneous and visceral fat pads, particularly in the dorsal shoulder and abdominal regions (Figure 3A). Both female and male NM1 KO mice consistently exhibited higher body weights compared to WT (Figure 3B-C). At 12 months, KO mice were 35% heavier on average. Abdominal adipose tissue accounted for approximately 29% of this difference and represented 11% of total body weight in KO mice compared to 4.6% in WT (Figure 3D-F). KO mice also had increased visceral adipose tissue and showed visually apparent obesity (Figure 4A). MicroCT imaging confirmed the progressive nature of adiposity in NM1 KO mice. At 2–4 months, KO and WT mice had similar thoracic and abdominal fat volumes (Figure 4A-B). By 12 and 18 months, KO mice displayed significantly greater adipose volumes in both regions (Figure 4B). At 12 months, KO thoracic adipose volume was 28.57 ±2.50 mm³ vs. 11.86 ±1.98 mm³ in WT (p<0.05), and abdominal volume was 20.69 ±2.60 mm³ vs. 8.59 ±1.13 mm³ in WT (p<0.05). These differences widened at 18 months, confirming an age-dependent adiposity phenotype (Figure 4B). To test whether increased fat accumulation could be attributed to hyperphagia, we monitored daily food intake in KO and WT mice over 8 days. Food consumption did not significantly differ between groups (Supplementary Figure 1H), suggesting that the obese phenotype in NM1 KO mice is not driven by increased caloric intake.

**Figure 3.**
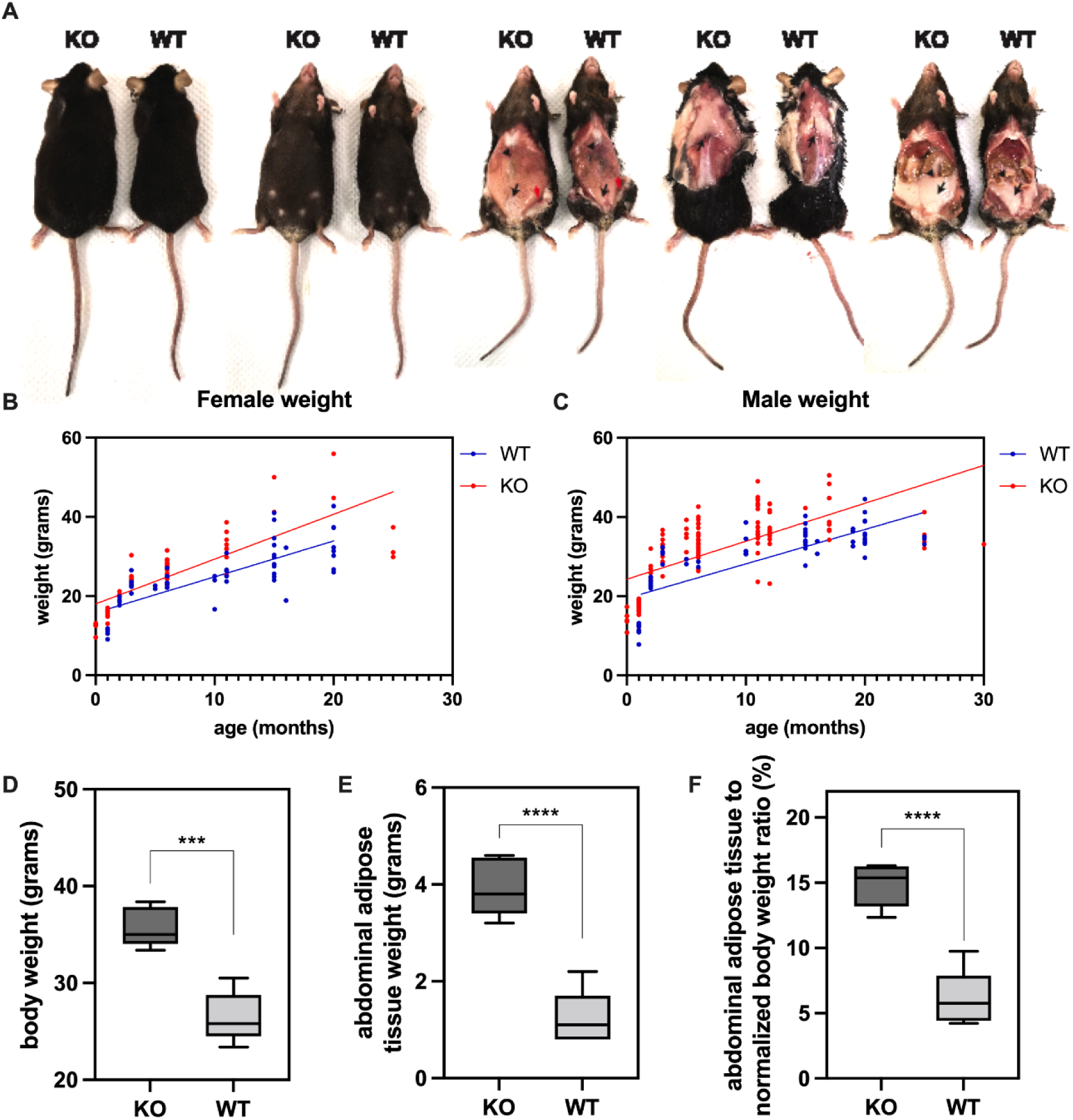
NM1 knockout leads to increased body weight and adipose tissue accumulation. (A) Representative images of WT and NM1 KO female mice, shown before and after dissection to visualize fat depots. KO mice display visibly larger subcutaneous and visceral fat pads, particularly in the dorsal shoulder and abdominal regions. (B–C) Longitudinal body weight measurements in female (B) and male (C) mice from 2 to 24 months of age. KO mice of both sexes gain more weight with age compared to WT controls. (D–F) Quantification of metabolic phenotypes in 12-month-old female mice. (D) Total body weight; (E) absolute weight of dissected abdominal adipose tissue; and (F) adipose tissue weight normalized to total body weight. KO mice display significantly greater adiposity across all measures. Data represents mean ± SEM. Statistical significance determined by unpaired t-test (***p < 0.001; ****p < 0.0001).

**Figure 4.**
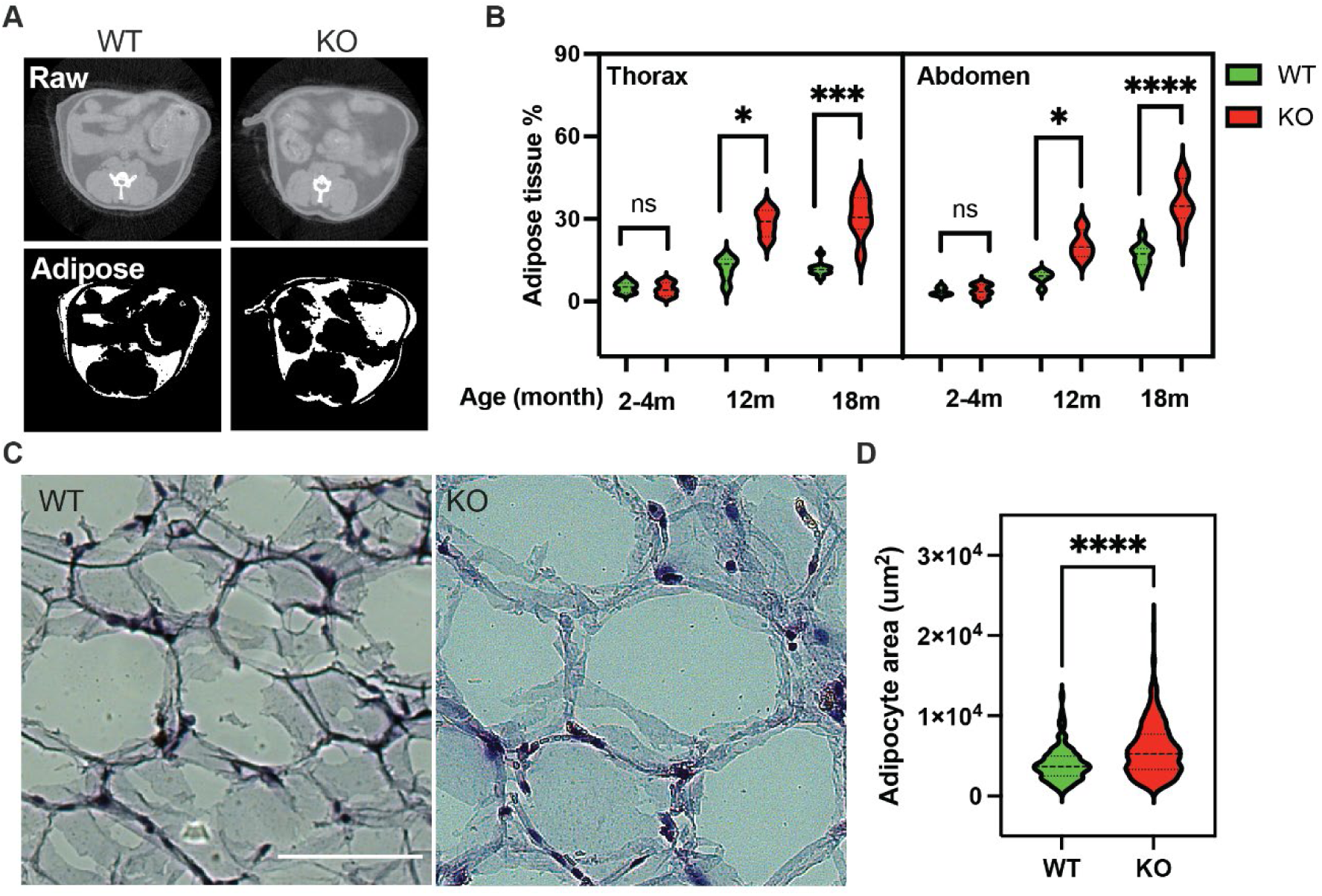
NM1 knockout increases adipose tissue accumulation and adipocyte hypertrophy *in vivo*. (A) Representative raw and segmented microCT images of the abdominal and thoracic regions from live wild-type (WT) and NM1 knockout (KO) mice, highlighting increased adipose tissue in KO mice. (B) Quantification of adipose tissue volume (% of total tissue) in the thoracic and abdominal regions at 2–4 months, 12 months, and 18 months of age. KO mice show a progressive and significant increase in fat accumulation with age. Each condition includes a minimum of four biological replicates. (C) Hematoxylin and Eosin (H&E) staining of visceral adipose tissue (VAT) sections from WT and KO mice at 18 months, showing adipocyte morphology. Scale bar: 100 μm. (D) Quantification of adipocyte surface area from H&E-stained sections confirms adipocyte hypertrophy in KO tissue. Data are presented as violin plots; statistical comparisons were made using unpaired t-tests (*p < 0.05; ***p < 0.001; ****p < 0.0001; ns, not significant).

Remarkably, histological analysis of abdominal VAT at 18 months revealed marked adipocyte hypertrophy in KO mice (Figure 4C). Mean adipocyte area was 5792 ±112.1 μm² in KO versus 3997 ±124.6 μm² in WT (p<0.0001). KO adipocytes spanned a wider range (733–22237 μm²) than WT (382–12731 μm²)(Figure 4D). These findings confirm that loss of NM1 leads to major morphological alterations characterized by hypertrophic expansion where existing adipocytes grow in size. This condition is less metabolically healthy and when adipose tissue expansion occurs primarily by hypertrophy, it often leads to dysfunctional adipose tissue with inflammation, hypoxia and, potentially, insulin resistance^33^.

Based on the above, we next studied the transcriptional changes that accompany NM1-dependent hypertrophy in the adipose tissue by bulk RNA-seq analysis on VAT isolated from WT and NM1 KO mice. Gene expression analysis of VAT revealed substantial transcriptional alterations in NM1 KO mice, with 1,142 genes significantly upregulated and 752 genes significantly downregulated compared to WT controls (Figure 5 A-B). To assess the biological significance of these changes, we performed GO enrichment analysis, focusing on biological processes (BP), cellular components (CC) and pathways (KEGG) (Figure 5C-E). Among the most enriched biological terms were the positive regulation of phosphatidylinositol 3-kinase/protein kinase B (PI3K/Akt) signaling, inflammatory response, and angiogenesis, suggesting disruptions in metabolic signaling, immune activation, and vascular remodeling. CC analysis revealed enrichment in extracellular matrix-related categories, including “collagen-containing extracellular matrix” and “extracellular region,” indicating potential structural changes in the VAT microenvironment. KEGG pathway analysis highlighted enrichment in endocrine resistance, PI3K/Akt signaling, and Rap1 signaling, further supporting a role for NM1 in regulating signal transduction and metabolic homeostasis, which is in agreement with previously published data^11^.

**Figure 5.**
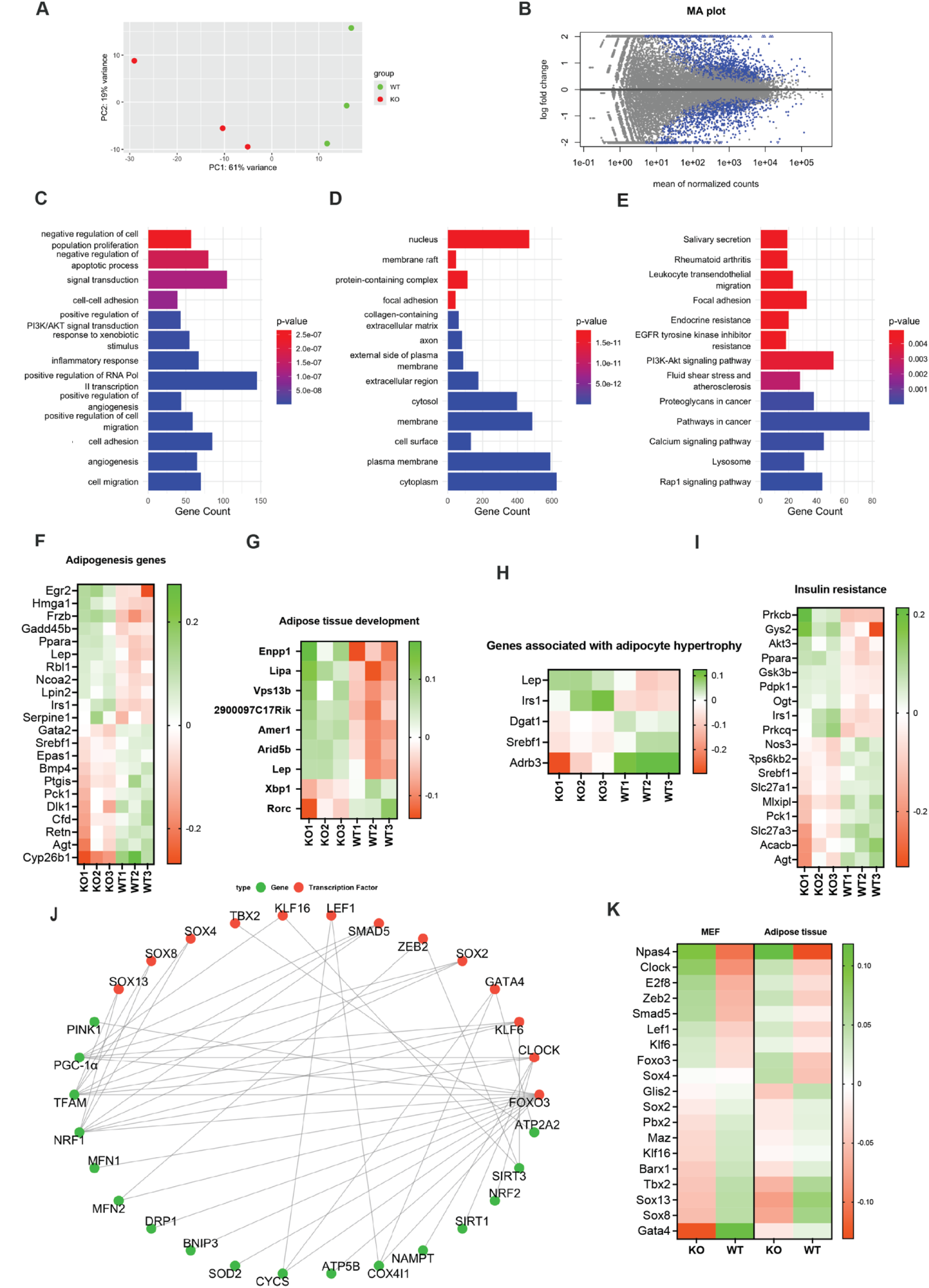
NM1 deletion alters transcriptional, mitochondrial, and adipogenic regulatory programs in visceral adipose tissue. (A) Principal component analysis (PCA) of RNA-seq data from wild-type (WT) and NM1 knockout (KO) visceral adipose tissue (VAT) samples, showing clear separation between genotypes. (B) MA plot displaying log fold change versus mean normalized gene expression. Blue dots represent significantly differentially expressed genes (adjusted p-value < 0.05). (C–E) Gene ontology (GO) and KEGG enrichment analysis of differentially expressed genes. (C) Biological process (BP), (D) cellular component (CC), and (E) KEGG pathway enrichment, with color indicating significance level. (F–H) Gene expression heatmaps of selected genes from RNA-seq. Each column represents an individual replicate. Color indicates increased (green) or decreased (red) expression in given sample. (F) Adipogenesis-related genes, (G) genes involved in adipose tissue development, and (H) genes associated with adipocyte hypertrophy. (J) genes associated with insulin resistance. Each column represents an individual replicate. (J) Literature-derived transcription factor–mitochondrial gene interaction network. Red nodes represent transcription factors; green nodes represent mitochondrial targets,. (K) Expression of the top 19 transcription factors (TFs) identified from regulatory network modeling (Figure 1) across MEFs and VAT.

To better understand changes related to adipocyte differentiation, we examined the expression of genes involved in adipogenesis (Figure 5 F-H). Several genes implicated in transcriptional regulation, lipid metabolism, and adipocyte function were significantly upregulated in KO VAT, including *Egr2, Hmga1, Frzb, Gadd45b, Ppara, Lep, Rbl1, Ncoa2, Lpin2, Irs1,* and *Serpine1*. However, key pro-adipogenic regulators such as *Gata2, Srebf1, Epas1, Bmp4, Ptgis, Pck1, Dlk1, Cfd, Retn, Agt,* and *Cyp26b1* were significantly downregulated, suggesting a disrupted adipogenic regulatory network. This transcriptional profile, while indicating partial activation of adipogenic pathways, is consistent with impaired adipocyte differentiation observed in KO-derived MSCs, which exhibited reduced capacity to differentiate into adipocytes *in vitro*. Despite this reduced differentiation potential, both MSC-derived adipocytes and VAT histology revealed enlarged adipocytes in KO samples, suggesting that adipocyte hypertrophy, rather than increased adipogenesis, may account for the observed gene expression changes.

Beyond adipogenesis, genes associated with adipose tissue development and remodeling were also differentially expressed. Upregulated genes included *Enpp1, Lipa, Vps13b, 2900097C17Rik, Amer1, Arid5b,* and *Lep,* many of which are linked to lipid metabolism, extracellular matrix reorganization, and adipocyte expansion. In contrast, *Xbp1* and *Rorc* genes involved in stress response and immune regulation were downregulated, suggesting compromised adipose tissue homeostasis in KO VAT.

Given the presence of hypertrophic adipocytes in KO VAT, we further analyzed genes associated with adipocyte hypertrophy (Figure 5H). Among the significantly upregulated genes, *Lep* and Irs1 stood out, suggesting increased leptin production and insulin signaling. Conversely, *Dgat1, Srebf1, and Adrb3* were downregulated, indicating potential impairments in triglyceride synthesis, lipid homeostasis, and β-adrenergic signaling. These results support the notion that adipocyte hypertrophy in KO VAT may arise from dysregulated lipid metabolism and altered insulin responsiveness. In addition, we observed widespread transcriptional alterations in genes associated with insulin resistance (Figure 5I). Notably, *Pparα, Irs1, Gsk3b, Prkcb*, and *Srebf1* were significantly downregulated in NM1 KO mice, consistent with impaired insulin signaling and lipid homeostasis. These changes, together with increased expression of *Agtr1* and *Prkcg*, indicate a pro-diabetic transcriptional shift in VAT. This transcriptional signature supports the hypothesis that NM1 loss contributes not only to adipocyte dysfunction but also to systemic insulin resistance.

To investigate the transcriptional mechanisms underlying these extensive gene expression alterations, we went back to single-cell RNA-seq from WT MEF and ATAC-seq data derived from NM1 WT and KO MEFs. Using bulk RNA-seq data from both MEFs and VAT, we assessed the expression levels of these 19 identified TFs, observing consistent dysregulation patterns across both tissues (Figure 5K). This conserved dysregulation between MEFs and VAT was further confirmed by integrative analysis, highlighting a systemic transcriptional reprogramming induced by NM1 deletion. Thus, the transcriptional changes observed at the single-cell level in MEFs also accurately reflect transcription factor dysregulation in VAT tissue.

To further explore the functional implications of these transcriptional regulators, we examined their roles in mitochondrial gene expression. Through literature mining and network-based analysis, we identified several differentially expressed TFs, most notably *Foxo3, Clock, Klf6*, and *Gata4* as known regulators of mitochondrial function. These factors influence the expression of nuclear-encoded mitochondrial genes involved in OXPHOS, mitophagy, and mitochondrial biogenesis, including *TFAM*, *Ppargc1a* (PGC-1α), *Sirt3*, *Nrf1*, *Drp1*, and *Pink1* (Figure 5J). Notably, Foxo3 has been shown to directly regulate *Pink1*, linking NM1 deletion to impaired mitophagy and mitochondrial quality control. Similarly, Clock modulates mitochondrial dynamics and respiration through regulation of *Drp1* and *Nrf1*. Network analysis of transcription factor– mitochondrial gene relationships revealed direct and indirect regulatory connections, supporting a model in which NM1 deletion disrupts mitochondrial homeostasis through transcriptional cascades. These changes underlie the altered energy metabolism and adipocyte dysfunction observed in KO VAT and are compatible with the morphological mitochondrial alterations and the metabolic reprograming from OXPHOS to aerobic glycolysis resulting from NM1 KO^11^.

Collectively, these findings support an essential role for NM1 in transcriptional regulation and metabolic reprograming during adipogenesis.

### Immune-driven transcriptional reprogramming in NM1-deficient visceral adipose tissue

To gain more mechanistic insights into NM1 dependent transcription regulation during adipogenesis, we next performed IPA of differentially expressed genes from NM1 KO VAT. The analysis revealed a pro-inflammatory regulatory network centered on *IFNG* and TNF (Figure 6E). These upstream regulators were predicted to activate multiple immune-related processes, including leukocyte migration, maturation of blood cells, and increased quantity of antigen-presenting cells^34^. Supporting this immune-driven response, downstream targets such as *IL33*, *TREM2*, and *CSF1* were also activated, while inhibitory signals involving *EIF4EBP1* and *GLIS2* were suppressed, indicating broad transcriptional remodeling.

**Figure 6.**
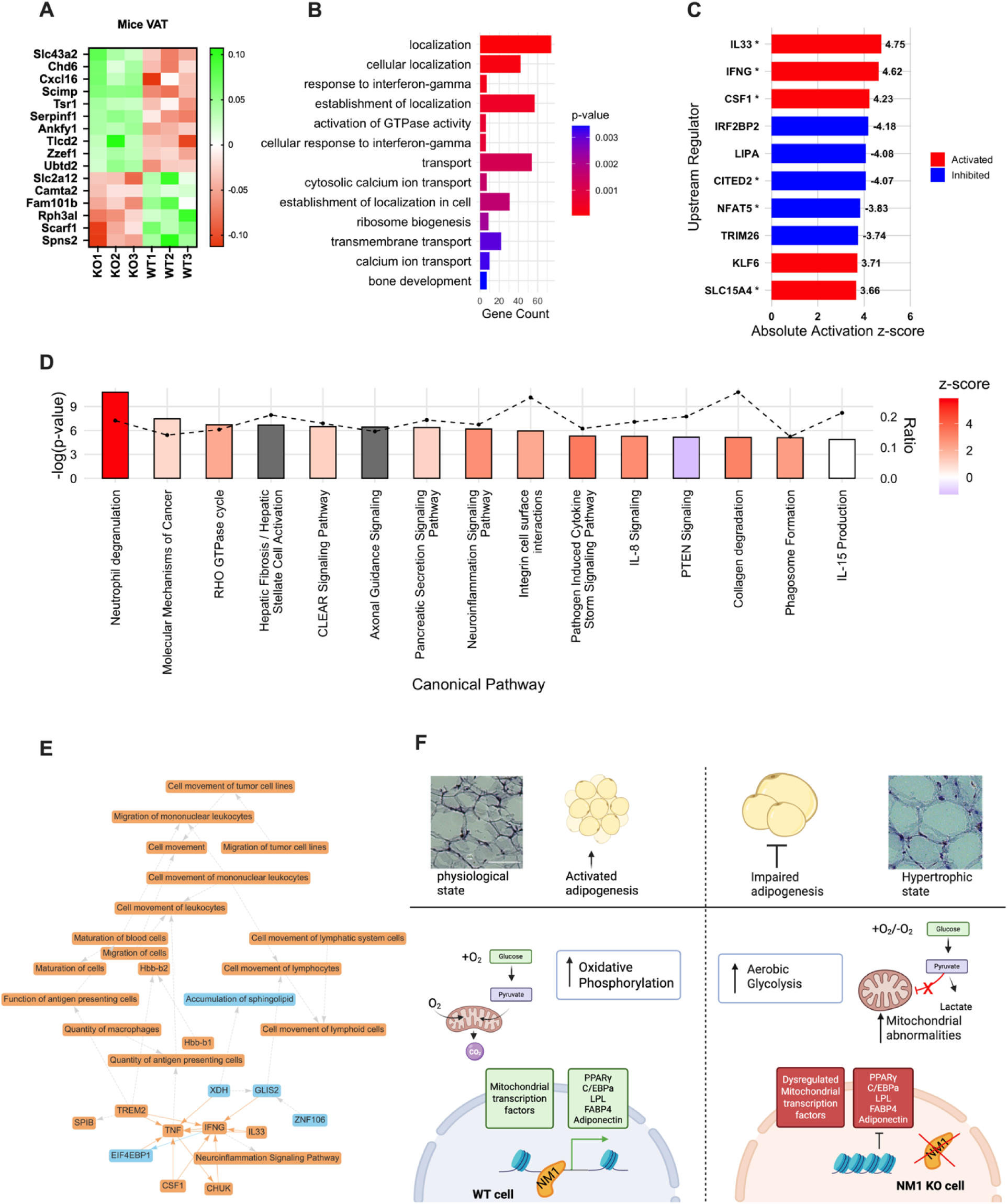
NM1 deletion remodels adipose transcriptional networks and reveals conserved regulatory links to human MYO1C-associated gene modules. (A)Heatmap showing the expression of the genes overlapping between mouse NM1 KO visceral adipose tissue (VAT) RNA-seq data and Community 184—a MYO1C-centered SNP-gene co-regulatory network derived from human VAT eQTL data (GTEx v8). Genes shown are significantly dysregulated in NM1 KO mice and are connected to key adipose-relevant SNPs. (B) Gene Ontology (GO) enrichment analysis of Community 184 using the topGO R package with the “elim” algorithm, highlighting biological processes enriched among overlapping genes. (C) Barplot showing the top 10 upstream regulators predicted by IPA, colored by activation z-score (red = activated, blue = inhibited). Asterisks (*) indicate regulators supported by IPA mechanistic network models. Values next to bars represent the signed activation z-score. These regulators represent high-confidence transcriptional drivers of the NM1-associated gene expression changes. (D) Canonical pathway enrichment analysis from IPA reveals significant activation of inflammatory and immune-related pathways. Bar colors reflect predicted activation state (z-score); dashed line shows significance threshold (−log p-value). (E) IPA graphical summary of upstream regulator and pathway activity predictions based on differentially expressed genes in NM1 KO VAT. Orange lines and nodes indicate activation, blue indicates inhibition. (F) Schematic summary of NM1’s role in adipocyte differentiation and mitochondrial regulation. In WT cells, NM1 supports chromatin accessibility at adipogenic and mitochondrial gene loci, promoting oxidative phosphorylation and adipocyte maturation. In contrast, NM1 deletion leads to transcriptional repression, impaired mitochondrial function, a shift toward aerobic glycolysis, and adipocyte hypertrophy.

Upstream regulator analysis (Figure 6C) identified *IL33*, *IFNG*, and CSF1 as the top predicted activated drivers (z-scores > 4), highlighting their central role in orchestrating inflammatory responses. In contrast, inhibited regulators such as *CITED2, NFAT5*, and *NPC1* may reflect suppression of transcriptional repression and metabolic control. The coordinated activation of immune-related cytokines alongside inhibition of regulatory and metabolic factors suggests a directional shift in adipose tissue identity^34^. These findings are consistent with the known roles of *IFNG* and *TNF* in adipose inflammation and insulin resistance^35,36^, and with the emerging function of *IL33* in type 2 immune remodeling of fat^37^.

Canonical pathway enrichment analysis (Figure 6D) further supported these findings, with Neuroinflammation Signaling, IL-17 Signaling, and Cell Movement of Lymphoid Cells ranking among the top pathways. The majority of the top 15 canonical pathways showed positive z-scores, indicating pathway activation consistent with enhanced leukocyte trafficking and cytokine signaling. Collectively, these IPA results suggest that NM1 deletion induces a transcriptional program characteristic of adipose tissue immune activation and inflammatory remodeling, aligning with the known immune-metabolic crosstalk observed in visceral fat depots^38^.

### NM1 KO in mouse VAT reveals regulatory links to human MYO1C and adipogenic networks

To investigate human regulatory architecture relevant to NM1 function, we examined Community 184, a SNP-gene co-regulatory module derived from human visceral adipose tissue using GTEx v8 eQTL data. SNP-gene interactions were filtered based on a loose statistical significance (FDR < 0. 20), a threshold chosen to limit the loss of connexons due to false negatives while limiting the noise brought by false positives^39^. They were then organized into distinct communities using modularity-based clustering algorithms. Community 184, which contains *MYO1C* (the closest human paralog to NM1), encompasses 224 genes and approximately 15,170 SNP-gene regulatory links spanning nearly every chromosome.

To explore the biological relevance of this regulatory module, we performed GO enrichment analysis using the topGO R package with the Îelim” algorithm, which accounts for hierarchical term structure and redundancy. This analysis revealed significant enrichment for biological processes associated with cytosolic transport (GO:0051179, p = 4.6e-5), GTPase activity (GO:0090630, p = 6.4e-4), and interferon-gamma signaling (GO:0034341, p = 2.8e-4), indicating functional convergence with transcriptional and immune-related pathways disrupted in NM1-deficient mouse VAT (Figure 6B).

To assess cross-species transcriptional conservation, we compared the NM1 KO mouse VAT RNA-seq dataset to Community 184. This analysis identified a subset of genes with significant overlap, which were also transcriptionally dysregulated in NM1 KO mice (Figure 6A). Heatmap visualization highlights altered expression of these conserved targets, several of which— including *Serpinf1*, and *Spns2* are linked to adipose tissue remodeling, immune signaling, and mitochondrial metabolism^40,41^.

To identify key regulatory variants within this network, we computed network centrality scores, including cores-scores that measure local centrality and outdegrees, the sum of edge weights going out of each SNP, that measure global centrality, to prioritize hub SNPs based on their connectivity to MYO1C and other highly co-regulated genes. Three SNPs, rs4968122, rs62068394, and rs8066936 emerged as central hubs within Community 184. These variants demonstrated strong eQTL effects in adipose tissue according to GTEx and were associated with adiposity- and metabolism-related traits in the NHGRI-EBI GWAS Catalog.

Functional annotations of the genes connected to these SNPs revealed involvement in lipid metabolism, cytoskeletal remodeling, and immune regulation, biological processes consistent with the known roles of NM1 and *MYO1C*. These cross-species parallels support an evolutionarily conserved role for the NM1/*MYO1C* family in orchestrating transcriptional networks that underlie adipose tissue function and metabolic homeostasis.

## Discussion

In this study, we uncover a new role for NM1 in coordinating adipogenic differentiation, mitochondrial, adipose tissue immune remodeling and homeostasis, revealing an unexpected regulatory layer linking genomic architecture with metabolic control. Our results identify NM1 as a central regulator of adipocyte differentiation, mitochondrial remodeling, and adipose tissue homeostasis. Using an integrative approach that combined epigenomics, transcriptomics, cell and tissue phenotyping, and cross-species network analysis, we demonstrate that NM1 acts at multiple regulatory layers to coordinate chromatin accessibility, transcription factor activity, and mitochondrial gene expression. These data reveal NM1 as a mechanistic link between nuclear architecture and metabolic reprogramming during adipogenesis.

NM1-deficient MSCs exhibit reduced adipocyte differentiation potential and gene expression, with a clear dosage-dependent defect across WT, HET, and KO genotypes. This is particularly notable for core adipogenic regulators such as *Pparg, Cebpa*, and *Adipoq*, whose expression is progressively reduced in HET and KO cells. The corresponding phenotype, fewer but hypertrophic adipocytes, parallels findings in other models of impaired adipogenesis^42,43^. In these settings, reduced differentiation capacity leads to lipid accumulation in fewer adipocytes, promoting hypertrophy and metabolic dysfunction.

Mechanistically, our integration of ATAC-seq and RNA-seq in NM1 KO MEFs identified a subset of genes whose differential expression was linked to changes in chromatin accessibility. Motif enrichment analysis revealed key adipogenic transcription factor motifs in differentially accessible regions, and network modeling identified 19 highly active transcription factors with consistent expression changes in both MEFs and VAT. This supports a direct role for NM1 in regulating gene expression through chromatin remodeling. Consistent with earlier work showing that NM1 interacts with the B-WICH complex to promote histone acetylation and transcriptional activation^35,36^, our findings extend its role to adipocyte-relevant transcriptional networks.

At the mitochondrial level, NM1 KO MSCs showed defective transcriptional activation of *TFAM* and *PINK1*, despite increased *PPARGC1A*. This decoupling is indicative of a block in mitochondrial biogenesis or turnover, which is essential for adipogenic differentiation and metabolic reprogramming^44,45^. These findings point to a functional impairment in the mitonuclear axis in the absence of NM1, potentially mediated through downstream transcriptional regulators such as *Foxo3* or *Clock*^46^, and reinforce NM1’s role in coordinating the nuclear transcriptional programs necessary for mitochondrial adaptation during adipogenesis.

*In vivo*, NM1 knockout mice developed progressive obesity with age, characterized by increased total and visceral fat mass and pronounced adipocyte hypertrophy. MicroCT scans revealed that KO mice accumulated fat rapidly after 12 months of age, and dissected fat pads confirmed greater adipose depot size. These changes were accompanied by significant transcriptional remodeling in VAT, including dysregulation of genes involved in lipid metabolism (*Dgat1, Lpin2*), insulin signaling (*Irs1, Lep*), and adipose tissue remodeling (*Enpp1, Arid5b*). The transcriptional profile in KO VAT suggests a partial activation of adipogenic programs alongside aberrant hypertrophic growth, which is typical of dysfunctional fat expansion^47^.

IPA analysis of VAT RNA-seq data revealed strong enrichment of pro-inflammatory signaling pathways, with predicted activation of *IFNG, TNF*, and *IL33*. These cytokines are implicated in adipose inflammation, insulin resistance, and the recruitment of immune cells into VAT^36,38^. The suppression of anti-inflammatory and metabolic regulators such as *CITED2* and *GLIS2* further supports a shift toward an immune-activated adipose tissue environment. Adipose inflammation is a well-known contributor to metabolic dysfunction, and the immune remodeling observed here may underlie some of the metabolic phenotypes seen in NM1 KO mice.

To investigate the conservation of NM1-associated regulatory networks, we analyzed a human adipose eQTL network centered on MYO1C, the closest paralog of NM1. Community 184 contained more than 15,000 SNP gene links and was enriched for genes involved in cytoskeletal regulation, interferon signaling, and intracellular transport. Several DEGs from mouse VAT, including *Cxcl16* and *Slc43a2*, were directly linked to adipose-specific regulatory SNPs in this network, and network centrality analysis prioritized rs4968122, rs62068394, and rs8066936 as potential human variants functionally linked to NM1/MYO1C function. The cross-species overlap between NM1-regulated mouse genes and human MYO1C eQTL targets suggests evolutionary conservation of a transcriptional module that regulates adipocyte biology and immune signaling. Given the increasing interest in nuclear actin and myosins in chromatin organization and transcriptional memory^48,49^, these findings raise the possibility that MYO1C genetic variation may influence obesity risk through similar chromatin-dependent pathways.

In summary, this study establishes NM1 as a chromatin-associated regulator that links transcription factor accessibility, mitochondrial biogenesis, and adipose tissue inflammation. NM1 deletion leads to defective adipocyte differentiation, mitochondrial dysfunction, VAT remodeling, and obesity, integrating nuclear mechanics with metabolic regulation. The convergence of mouse NM1 deletion and human *MYO1C* network topology suggests that this actomyosin pathway may represent a conserved regulatory axis in adipose biology, warranting further exploration in metabolic disease models and patient populations. NM1 and *MYO1C* therefore represent promising targets for further investigation in the context of obesity and type 2 diabetes, with potential for translational applications that bridge nuclear remodeling and metabolic homeostasis (Figure 6F).

## Methods

### Single-Cell preprocessing and filtering

Raw 10x Genomics data (accession: GSE264266_RAW)^50^ were downloaded from the Gene Expression Omnibus (GEO). Briefly, the dataset was quality-controlled, normalized, and clustered to ensure that only cells exhibiting canonical expression profiles consistent with homeostatic MEFs were retained for downstream analysis. Initial quality control involved filtering out cells expressing less than 200 genes. Genes detected in fewer than 10 cells were removed. Cells with a mitochondrial gene content greater than 15% were excluded. Each cell’s total counts were normalized to a target sum of 10000, and log1p transformation was then applied. Principal component analysis (PCA) was employed to reduce the dimensionality of the data and build a nearest neighbor graph, which served as the basis for the UMAP. This resulted in a dataset of 6869 cells by 16949 genes.

### Bulk RNA preprocessing and filtering

Raw count data from the GSE133506 dataset were merged into a count matrix and annotated with sample metadata (genotype and replicate) as well as gene symbols. The dataset was then processed using DESeq2 for normalization, dispersion estimation, and filtering of lowly expressed genes. Surrogate variable analysis (SVA) was performed to correct for hidden confounders by incorporating 1 surrogate variable (SV1) into the model, which also included genotype and replicate. Differential expression analysis was conducted using the apeglm shrinkage estimator, and results were visualized with an Enhanced Volcano plot and a heatmap derived from variance-stabilized data to assess sample clustering.

Similarly raw count data from six adipose tissue samples (3 KO and 3 WT) were merged into a single count matrix and annotated with genotype and replicate metadata, then subjected to the same pipeline, incorporating two surrogate variables (SV1 and SV2) into the model. Finally, the differential expression results of both analyses were saved to their respective CSV files (Code available at https://github.com/NYUAD-Core-Bioinformatics/Khalaji-et-al-2025/). Data are available via GEO (GSE236679 and GSE133506) and Supplementary table1.

### Bulk ATAC–seq data processing

ATAC-seq libraries were obtained from NCBI’s GEO (Data are available via GEO GSE198988) and processed following the protocol described previously^11^. Briefly, adapter sequences were removed using cutadapt, and the trimmed reads were aligned to GRCm38 using BWA. Reads with mapping quality scores below Q30 were discarded, duplicates were removed with Picard, and mitochondrial as well as unmapped reads were excluded. Peaks were then called separately for the WT and KO conditions using MACS3 callpeak, merged across conditions with bedops, and compiled into a peak-by-sample count matrix using subreads featureCounts, the final set of peaks was annotated using HOMER. Normalization and differential accessibility were assessed using edgeR.

### Bulk ATAC and RNA integration

Differential expression analysis between WT and KO NM1 MEFs was conducted by selecting genes with an adjusted p-value ≤ 0.05 and an absolute log2 fold change (|logFC|) ≥ 0.25. These genes were then linked to differentially accessible chromatin regions that met the criteria of FDR ≤ 0.05 and |logFC| ≥ 0.1 (see Figure1A). Each accessible region was subsequently analyzed using FIMO to identify transcription factor motif enrichment, employing the HOCOMOCO transcription factor database.

### Network inference

Gene regulatory network inference was performed using a single-task learning algorithm implemented in the Inferelator pipeline (https://github.com/flatironinstitute/inferelator). In brief, single RNA seq count matrix (cells × genes) was modeled as the product of a transcription factor activity (TFA) matrix (cells × transcription factors) and a regulatory network matrix (transcription factors × genes). First, a prior network was constructed by integrating chromatin accessibility data to identify transcription factor binding motifs and link them to nearby genes, using the inferelator-prior tool with the HOCOMOCO TF database. The TFA matrix was then computed by inverting this linear system. Next, both the gene expression data and the estimated TFA were input into the single-task BBSR pipeline to infer regulatory interactions, which were visualized using PyVis. To simulate NM1 knockout expression, TFA from the bulk RNA KO experiment was multiplied by the wild-type regulatory network, and the simulated expression values were compared to the observed measurements (R² = 0.65) to evaluate network accuracy. Scripts for analysis.

### Hi-C library preparation and analysis

Hi-C libraries were prepared from WT and NM1 KO MEFs as previously described (GEO accession GSE198989)^11^. Reads were processed using HiCUP with Arima-specific parameters, and compartment switching was analyzed using HOMER’s runHiCpca.pl (resolution = 500 kb). Differential compartments were defined as regions with significant PC1 polarity changes (FDR < 0.05). TAD boundaries and insulation scores were computed using findTADsAndLoops.pl and differential scores assessed with getDiffExpression.pl.

### Isolation of mesenchymal stem cells from compact bone

MSCs were isolated from compact bone of NM1 WT and KO mice aged 5–8 weeks. Mice were anesthetized and euthanized via cervical dislocation in accordance with institutional ethical guidelines. Femurs and tibias were collected under sterile conditions, and soft tissue was carefully removed. Bones were rinsed in phosphate-buffered saline (PBS), and bone marrow was flushed out using a 27G needle. The remaining bone shafts were crushed into 2–3 mm fragments and digested in Collagenase Type II (1 mg/mL in α-MEM) at 37°C for 1 hour with continuous rotation. Digested fragments were washed three times with PBS and cultured in α-MEM (without nucleotides), supplemented with 20% fetal bovine serum (FBS) and 1% penicillin/streptomycin. Cells were incubated under hypoxic conditions (0% oxygen, 5% CO₂) at 37°C to promote MSC migration from the bone matrix. Media was refreshed every 3 days, and adherent cells were harvested after 10–15 days using 0.05% trypsin-EDTA for expansion^51^.

### Adipogenic differentiation of mesenchymal stem cells

MSCs were seeded into 12-well plates and allowed to reach ∼90% confluency within 24 hours. Adipogenic differentiation was induced by supplementing α-MEM with 20% FBS, 1 μM dexamethasone, 10 μg/mL insulin, 1 μM rosiglitazone, and 0.5 mM IBMX. Cells were maintained in adipogenic medium for 20 days, with media changes every 3 days. Negative control cells were cultured under the same conditions in α-MEM supplemented only with 20% FBS.

### Quantitative RT-PCR analysis

Total RNA was extracted from MSCs collected at day 0 and day 20 of adipogenic differentiation using RNazol (Sigma-Aldrich). RNA purity and concentration were evaluated using NanoDrop and Qubit fluorometry. cDNA synthesis was carried out using the RevertAid First Strand cDNA Synthesis Kit (ThermoFisher Scientific) with 1 μg of RNA per reaction. qPCR was performed using Maxima SYBR Green qPCR Master Mix on a StepOnePlus Real-Time PCR System. Each sample was analyzed in four technical replicates, and expression levels were normalized to WT day 0 using GAPDH as the reference gene. Gene expression data represent the mean of three biological replicates. Primer sequences are listed in Supplementary table1.

### Animal Studies

All animal experiments were conducted in accordance with protocols approved by the Institutional Animal Care and Use Committee (IACUC) at New York University (approved protocol number: 23-0009A1). Mice were housed under Specific Pathogen-Free (SPF) conditions in standard individually ventilated cages, maintained on a 12-hour light/dark cycle, and provided ad libitum access to standard chow and water.

The NM1 knockout (NM1 KO) mice, on a C57BL/6 genetic background, were generated and characterized as previously described^20^. Age- and sex-matched C57BL/6 WT mice were purchased from The Jackson Laboratory and used as controls in all experiments.

### Mouse body weight measurement and adipose tissue dissection

Body weight of mice was recorded across the population of WT and NM1 KO mice during ageing using a calibrated digital scale. For the adipose tissue volume measurements, 12 months old recorded in WT (n=5) and NM1 KO (n=5) mice during aging IACUC protocols via cervical dislocation. Abdominal and anterior subcutaneous adipose tissues were dissected and weighed using an analytical balance. Data were normalized to total body weight.

### MicroCT imaging of adipose tissue

*In vivo* microCT scans were performed at 2–4 months, 12 months, and 18 months of age on WT and NM1 KO mice using the SkyScan 1276 system (Bruker, USA). Mice were anesthetized with 2.5% isoflurane and positioned in dorsal recumbency for imaging. Scanning parameters included 75 μm resolution, 50 kV, 200 μA, and 0.5 mm Al filter. Reconstructions were completed using NRecon with 20% beam hardening and level 2 ring artifact correction. 3D alignment was done in DataViewer. Adipose tissue segmentation and quantification (thoracic and abdominal) were carried out in CTAn using global thresholding and morphological filtering.

### Histological analysis and adipocyte quantification

VAT was collected from 18-month-old mice (n=3 per genotype), fixed in 4% PFA at 4°C for 24 h, and cryoprotected in 30% sucrose for 48 h. OCT-embedded samples were sectioned at 35 nm using a Leica cryostat. Hematoxylin and eosin (H&E) staining was performed following standard protocols^23^. Images were acquired using a Leica DMI6000 widefield microscope. Adipocyte surface area and density were quantified using ImageJ with the Adiposoft plugin. Calibration was performed based on microscope settings, and only intact cells were included. Each frame represented 0.8 mm². Statistical comparisons were made using GraphPad Prism with unpaired t-tests.

### RNA-Seq and IPA analysis

VAT RNA from 18-month-old WT and KO mice was extracted using RNAzol and homogenized with a Bead Ruptor 96. Libraries were prepared with NEBNext Ultra II RNA Library Prep Kit and sequenced on Illumina NextSeq 500/550. Reads were trimmed with Trimmomatic and quality-checked with FastQC. Alignment to GRCm38 was done with HISAT2, and gene counts were generated with HTSeq-count. Data were processed in NASQAR for normalization and differential expression. DEGs were defined as |log2FC| ≥ 1 and adj. p < 0.05. Enrichment was performed with Database for Annotation, Visualization, and Integrated Discovery (DAVID). Data are available via GEO (GSE236679 and GSE133506) and Supplementary table1.

Differential expression data were analyzed using Ingenuity Pathway Analysis (IPA, QIAGEN). Genes with |log2FC| ≥ 0.5 and adj. p < 0.05 were used. Canonical pathways, upstream regulators, and functional networks were assessed using z-score algorithms.

### SNP network and cross-species annotation

eQTL networks in human VAT (GTEx v8) were used obtained from the study from Gaynor et al.^53^, and SNP-gene modules were identified using a bipartite modularity maximization algorithm^54^ implemented in the bioconductor netZooR package^55^. Community 184, containing MYO1C enriched for adipose-relevant SNPs and GO terms (the R bioconductor topGO package v. 2.44, using the elim method and all the genes of the eQT networks as background, see Stone et al.,^55^), was extracted for further analysis. Network summary statistics measuring SNP connectivity were obtained from study by Stone et al.,.^55^). Top SNPs were prioritized based on connectivity, GTEx significance, and GWAS associations. Cross-species gene functions were annotated using GeneCards, NCBI, and PubMed.

## Acknowledgments

This work was supported by grants from New York University Abu Dhabi, by Tamkeen under the NYU Abu Dhabi Research Institute Award to the NYUAD Center for Genomics and Systems Biology (ADHPG-CGSB), and by the Sheikh Hamdan Bin Rashid Al Maktoum Award for Medical Sciences to PP. We also acknowledge the institutional support from the Institute of Molecular Genetics of the Czech Academy of Sciences (RVO: 68378050) and the Research Program Strategy AV21 Future of Assisted Reproduction (ART) (AV21-VP38/2025) provided by the Czech Agency of Sciences to TV. RNA sequencing was performed by the NYUAD CGSB Core at New York University Abu Dhabi and Bioinformatics assistance was provided by the NYUAD Bioinformatics Core at New York University Abu Dhabi. This research was partially carried out using the Core Technology Platforms resources at New York University Abu Dhabi. We would like to thank Marc Arnoux for assistance with deep sequencing and Nizar Drou for assistance with the analysis.

## Author Contributions

SKh performed the majority of experiments, analyzed the sequencing data, and wrote the manuscript together with PP. TV conceptualized the research project, performed initial phenotyping and contributed to manuscript writing. SKh, VF, RS and SKa set up initial mouse dissection experiments. ZL was involved in the histology of adipose tissue. MB contributed to the microCT scan experiments and quantification of adipose tissue. GS and MF performed the bioinformatics analysis. PP supervised the research project, wrote the manuscript, and analyzed the data.

## Conflict of interest

There is no conflict of interest.

## Notes

https://www.ncbi.nlm.nih.gov/geo/

### Competing Interest Statement

The authors have declared no competing interest.

