## Supplemental material for "Nuclear Myosin 1 links genomic architecture to adipose tissue remodeling, metabolic inflammation and obesity in mice"

**Supplementary material**

**Supplementary Figure 1. NM1 regulates adipogenic transcriptional networks and metabolic programming independent of food intake.** (A) Observed versus predicted gene expression values for all genes included in the transcription factor activity (TFA) model. The red dotted line indicates the ideal fit ($R^2 = 0.63$; Pearson $r = 0.80$), demonstrating the predictive strength of the inferred network. (B) Transcription factor–target gene regulatory network inferred by integrating single-cell RNA-seq and bulk RNA-seq data using the Inferelator framework. Red and blue edges indicate predicted positive and negative regulatory interactions, respectively. Green nodes represent target genes; red nodes represent transcription factors. (C–G) Heatmaps showing gene expression patterns from RNA-seq of visceral adipose tissue (VAT) across WT and NM1 KO mice, grouped by functional pathway. Panel (C) shows genes involved in insulin signaling; (D) insulin secretion; (E) negative regulation of fat cell differentiation; (F) positive regulation of fat cell differentiation; and (G) endocrine resistance. Expression values are scaled as z-scores across samples. (H) Daily food consumption measured over 8 days in individually housed WT ( green) and NM1 KO (red) female mice (n = 3 per group). No significant differences in food intake were observed between groups across the measurement period. (I) Body weight measurements of the same mice taken daily over 9 days.

### Supplementary figure 1

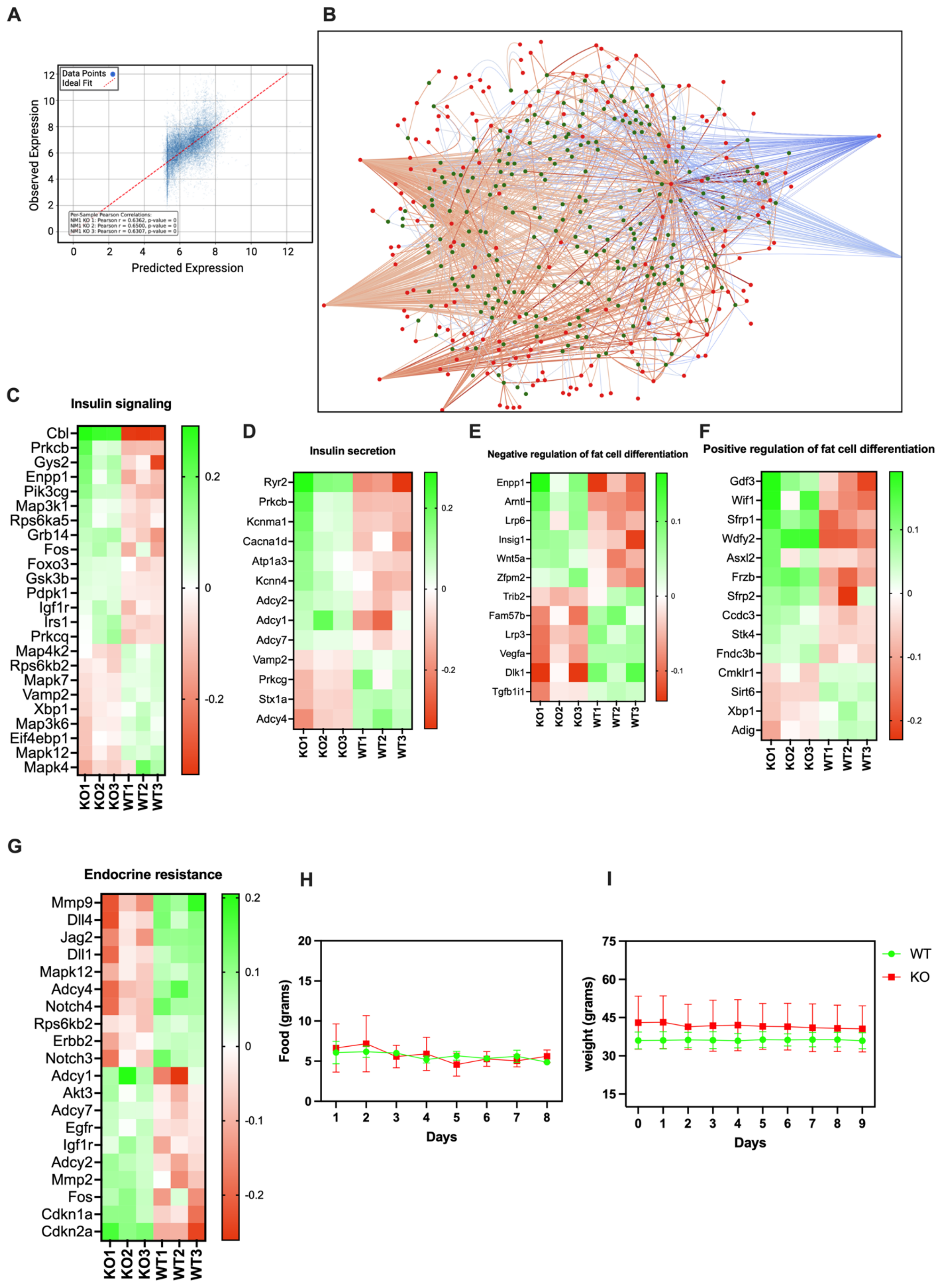

**Supplementary table 1. deseq (bulk RNA seq) file from adipose tissue**

**Supplementary table 2. Gene list from predictive model ATAC-seq vs bulk RNA-seq**

**Supplementary table 3. Results from MYO1C eQTL network**

**Supplementary Table 4. The primer sequences used in qRT-PCR experiments.**

| Gene name | Sequences |
|---|---|
| Pparg_F | TGTTATGGGTGAAACTCTGGG |
| Pparg_R | AGAGCTGATTCCGAAGTTGG |
| Adipoq_F | TGTCTGTACGATTGTCAGTGG |
| Adipoq_R | GCAGGATTAAGAGGAACAGGAG |
| Fabp4_F | GACAGGAAGGTGAAGAGCATC |
| Fabp4_R | GTCACGCCTTTCATAACACATTC |
| LPL_F | AACAAGGTCAGAGCCAAGAG |
| LPL_R | CCATCCTCAGTCCCAGAAAAG |
| Cebpa_F | CATGCCGGGAGAACTCTAAC |
| Cebpa_R | CTGGAGGTGACTGCTCATC |
| Cebpb_F | GTTTCGGGACTTGATGCAATC |
| Cebpb_R | TTTAAGGTGATTACTCAGGGCC |
| Ppargc1_ F | CACCAAACCCACAGAAAACAG |
| Ppargc1a_ R | GGGTCAGAGGAAGAGATAAAGTTG |
| TFAM_F | CACCCAGATGCAAAACTTTCAG |
| TFAM_R | CTGCTCTTTATACTTGCTCACAG |
| GAPDH_F | CATCTTCCAGGAGCGAGACC |
| GAPDH_R | CCTTCAAGTGGGCCCCG |
| PINK1_F | GACTCCCACCTTTCCCTTTG |
| PINK1_R | GTAACTGCTCCATACTCTCCAG |
